# A brief glimpse at a haptic target is sufficient for multisensory integration in reaching movements

**DOI:** 10.1101/2020.10.04.325712

**Authors:** Ivan Camponogara, Robert Volcic

**Author notes:** Contact information: Ivan Camponogara, Department of Psychology, New York University Abu Dhabi, PO Box 129188, Abu Dhabi, United Arab Emirates.

## Abstract

Goal-directed aiming movements toward visuo-haptic targets (i.e., seen and handheld targets) are generally more precise than those toward visual only or haptic only targets. This multisensory advantage stems from a continuous inflow of haptic and visual target information during the movement planning and execution phases. However, in everyday life, multisensory movements often occur without the support of continuous visual information. Here we investigated whether and to what extent limiting visual information to the initial stage of the action still leads to a multisensory advantage. Participants were asked to reach a handheld target while vision was briefly provided during the movement planning phase (50 ms, 100 ms, 200 ms of vision before movement onset), or during the planning and early execution phases (400 ms of vision), or during the entire movement. Additional conditions were performed in which only haptic target information was provided, or, only vision was provided either briefly (50 ms, 100 ms, 200 ms, 400 ms) or throughout the entire movement. Results showed that 50 ms of vision before movement onset were sufficient to trigger a direction-specific visuo-haptic integration process that increased endpoint precision. We conclude that, when a continuous support of vision is not available, endpoint precision is determined by the less recent, but most reliable multisensory information rather than by the latest unisensory (haptic) inputs.

## 1 Introduction

Everyday movements are not under exclusive visual control, but are often supported by the simultaneous use of visual and haptic—proprioceptive and tactile—inputs (Camponogara and Volcic, 2019b,a, 2021). For instance, when tapping on our phone, visual information is combined with the proprioceptive and tactile information from the hand holding the phone to guide contralateral hand movements. These actions toward multisensory targets are usually characterized by a higher precision than those toward unisensory—visual only or haptic only—targets (Desmurget et al, 1997; van Beers et al, 1999b,a; Monaco et al, 2010; van Atteveldt et al, 2014; Cameron and López-Moliner, 2015). This higher precision is achieved by the simultaneous availability of visual and haptic information both during the action planning phase (i.e., before the movement onset) and during movement execution.

Studies on visuomotor and multisensory-motor integration have shown that providing vision during movement execution is a *conditio sine qua non* to guarantee the best movement performance. Specifically, reaching movements when vision is available during both movement planning and movement execution are more precise than when vision is withheld just after movement onset (Keele and Posner, 1968; Elliott and Madalena, 1987; Blouin et al, 1993; Rossetti et al, 1994; Westwood et al, 2001; Khan et al, 2002, 2006; Kennedy et al, 2015; Tremblay et al, 2017). Similarly, the precision of reaching movements toward visuo-haptic targets is also reduced if vision is provided only during the planning phase, even though haptic information is provided for the whole movement duration (Desmurget et al, 1997; Monaco et al, 2010; Cameron and López-Moliner, 2015). Importantly, multisensory planned actions are still more precise than when only vision is provided during the planning phase, or, when only haptic target information is available for the whole action execution. This suggests that a brief visual exposure before movement onset is sufficient for initiating a multisensory integration process that leads to an advantage that persists throughout the whole action.

However, it is not yet clear (a) whether the duration of visual exposure directly affects movement precision, and, (b) if there is a critical visual exposure duration needed for the multisensory advantage to occur. Studies performed so far have provided simultaneous visual and haptic target information before movement onset for a substantially long time window which spanned at least two seconds of the movement planning phase, without systematically exploring the role of vision in the planning of multisensory reaching movements (Desmurget et al, 1997; Monaco et al, 2010; Cameron and López-Moliner, 2015; Khanafer and Cressman, 2014).

To investigate whether and how the modulation of the visual exposure duration during the planning phase affects action performance, we performed an experiment in which participants underwent five separate blocks of trials. In a Hapto-Visual block (HV_t_), participants were haptically sensing the object with their left hand while very brief vision (50 ms, 100 ms, 200 ms or 400 ms) was provided following a start signal. In these cases, multisensory target information was accessible only during the planning phase (50 ms, 100 ms, 200 ms visual exposures) or partially also during the early movement execution phase (longest visual exposure, 400 ms). In an additional Visual block (V_t_), the same visual exposure durations were provided, but without any additional haptic feedback about the target position. Thus, in the V_t_ block, participants had only a short time interval to visually locate the object before performing the action without any other visual or haptic feedback. Three more blocks of trials were run in which: participants were holding the object with their left hand, but vision was prevented throughout the whole movement (Haptic Full: H_Full_), only visual target information was available for the whole movement (Visual Full: V_Full_), or, visual and haptic information was available for the entire movement (Hapto-Visual Full: HV_Full_).

Based on previous studies (van Beers et al, 1999b,a), we expect a higher precision in the multisensory block compared to each unisensory block. Moreover, we anticipate a decrease in precision when multi-sensory information is not provided during action execution, but a better performance compared to when only haptics is available, as reported by Monaco et al (2010) and Cameron and López-Moliner (2015). Additionally, if a critical visual exposure duration is needed for multisensory integration, we expect the multisensory advantage to be limited to only specific visual exposure durations. On the other hand, if even a 50 ms glimpse at the haptic target is sufficient to trigger a multisensory integration process, we expect an improved performance compared to the unisensory haptic and visual conditions at all visual exposure durations.

## 2 Method

### 2.1 Participants

Thirteen students from the New York University Abu Dhabi took part in this study (1 male, age 19.3 ± 0.48). All had normal or corrected-to-normal vision and no known history of neurological disorders. All of the participant were naïve to the purpose of the experiment and were provided with a subsistence allowance. The experiment was undertaken with the understanding and informed written consent of each participants and experimental procedures were approved by the Institutional Review Board of New York University Abu Dhabi in compliance with the Code of Ethical Principles for Medical Research Involving Human Subjects of the World Medical Association (Declaration of Helsinki).

### 2.2 Apparatus

The stimulus consisted of a 3D printed cylinder with a diameter of 35 mm, and a height of 50 mm. A second cylinder with a diameter of 10 mm and a height of 50 mm was used as the home position (Figure 1 A). To avoid any learning effect, the stimulus object was randomly presented during the experiment either at 150 mm or at 300 mm distance from the home position in the sagittal direction (depth). The 150 mm distance was used on a smaller proportion of trials to make the target position less predictable. These data were not used in the analysis.

**Figure 1:**
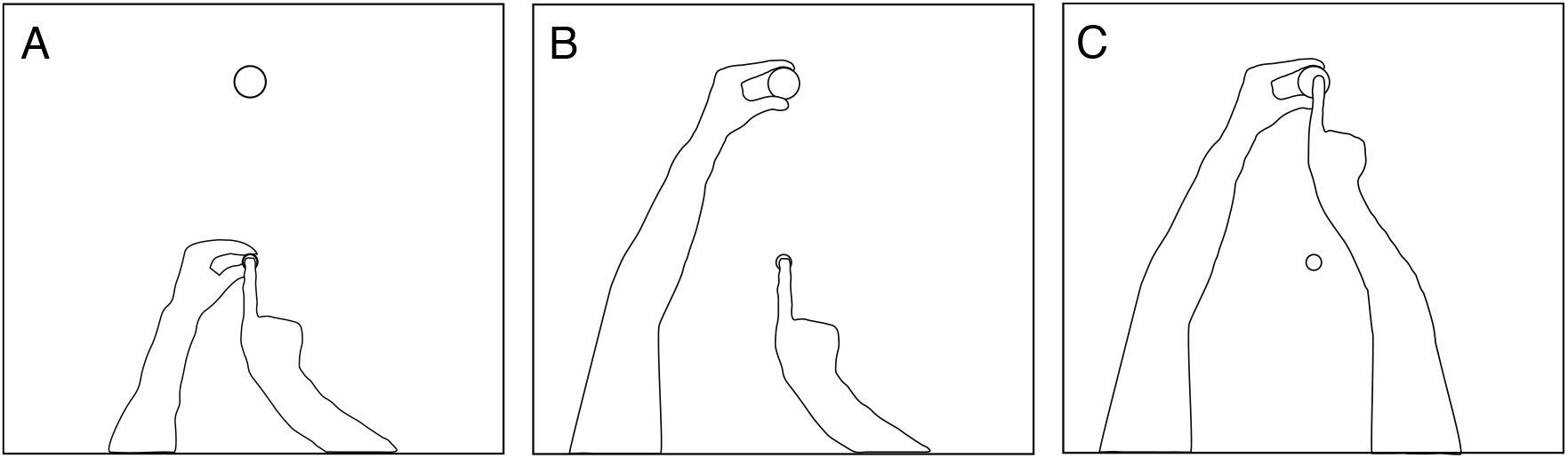
Top view of the experimental setup and task. A) All the trials started with the index digit of the right hand positioned at the top of the home position and the left hand holding the basis of the home cylinder. B) In the HV_t_, H_Full_, and VH_Full_ blocks, participants were asked to touch and hold the target object with their left hand (two-digit grasp) before each trial. C) After the start tone was delivered, participants reached for the top of the target cylinder with their right hand. The trial sequence in V_t_ and V_Full_ blocks was the same except that participants kept their left hand at the home position.

A pair of occlusion goggles was used to control vision availability during the trials (Red Scientific, Salt Lake City, UT, USA). A pure 1000 Hz tone, 100 ms in length, was used to signal the start of the trial, whereas a 600 Hz tone with the same length was used to signal its end.

Index movement was acquired on-line at 200 Hz with a sub-millimeter resolution by using an Optotrak Certus system (Northern Digital Inc., Waterloo, Ontario, Canada). The position of the fingertip was calculated during the system calibration phase with respect to three infrared-emitting diodes attached to the distal phalanx that acted as a rigid body (Nicolini et al, 2014). By tracking this rigid body we were then able to determine the exact position of the fingertip during the experimental sessions. The Optotrak system was controlled by the MOTOM toolbox (Derzsi and Volcic, 2018) and the occlusion goggles were controlled by a custom Matlab program.

### 2.3 Procedure

Participants sat comfortably in front of a table with their torso touching its edge. All the trials started with the participants’ index digit of the right hand positioned at the top of the home position, the left hand positioned at the basis of the home cylinder and the occlusion goggles closed (Figure 1A). Before each trial, the target cylinder was positioned at 300 mm or 150 mm. Participants were required to perform a rapid and accurate right-hand reaching movement to the center of the target cylinder.

In a first session, we performed a Hapto-Visual (HV_t_) and a Visual (V_t_) block, where we manipulated the availability of visual information along the movement. In both blocks, vision was provided for 50 ms, 100 ms, 200 ms or 400 ms following the start tone (Figure 2, first four rows). In the HV_t_ block, participants were requested to touch the object with the left hand and hold it with their index and the thumb from the beginning of each trial before the start tone was delivered (Figure 1B). In the V_t_ block, instead, participants were requested to keep the left hand at the home position for the whole duration of the trial. Therefore, while in the HV_t_ block participants could use both visual and haptic information about the position of the stimulus object, in the V_t_ block participants had only few milliseconds to detect the position of the object and integrate this information into the action plan.

**Figure 2:**
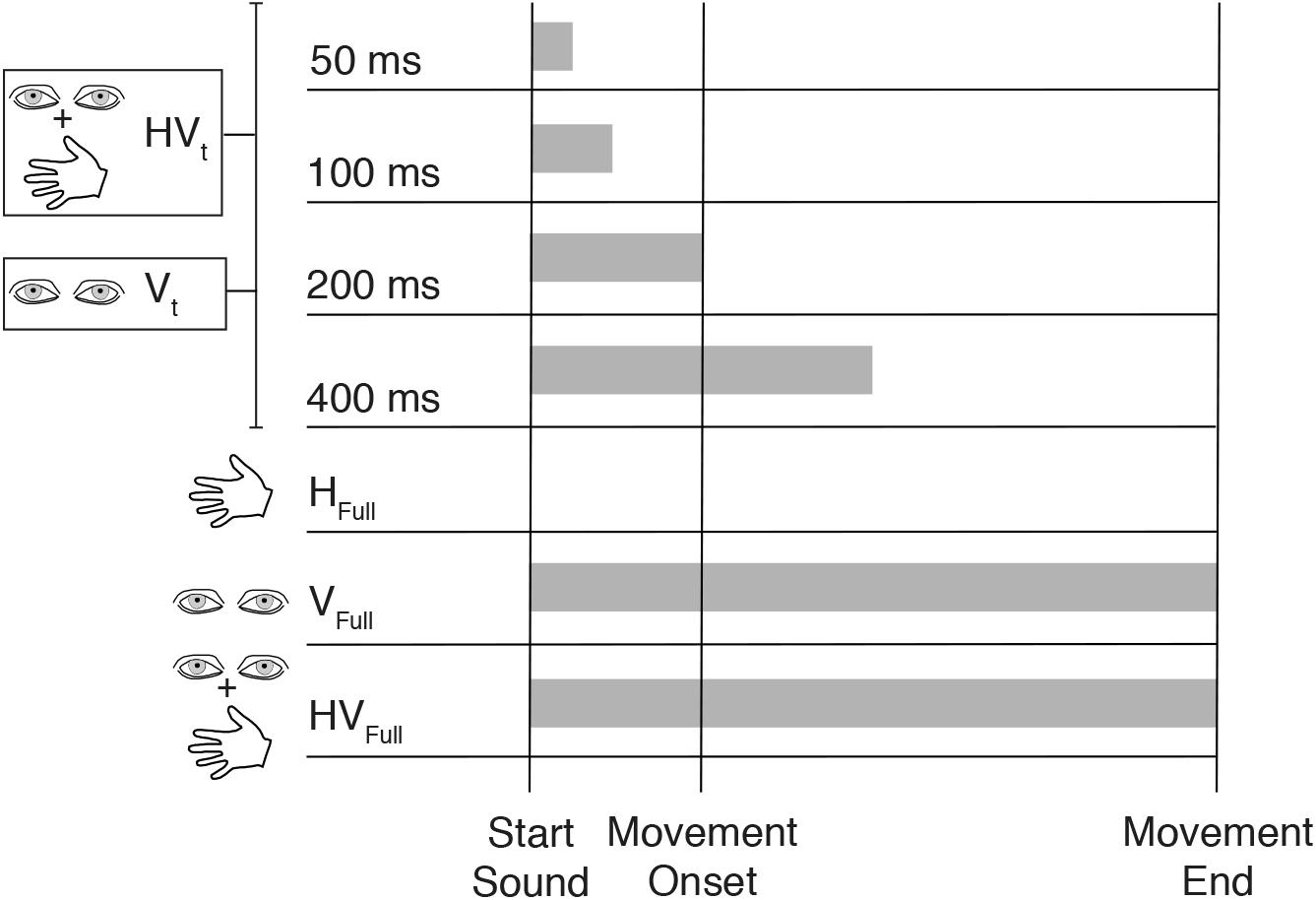
Schematic representation of the experimental conditions. The grey horizontal bars represent the period during which vision was available. The first four rows represent the conditions performed in H_Vt_ and V_t_ blocks. Participants started each trial with the goggles closed and vision randomly provided for 50 ms, 100 ms, 200 ms, or 400 ms. In the H_Vt_ block participants had concurrent haptic object information provided by the left hand. The last three rows represent the conditions performed in H_Full_, V_Full_ and HV_Full_ blocks. In the H_Full_ block, vision was prevented and participants were feeling the object with their left hand. In V_Full_ and HV_Full_ blocks, vision was allowed for the whole duration of the trial, which was supplemented by the haptic information in the HV_Full_ block.

In a second experimental session, we performed the H_Full_, V_Full_ and HV_Full_ blocks. In the H_Full_ block, participants were asked to touch and hold the object with the left hand from the beginning of each trial, while the goggles remained closed for the whole trial. Thus, participants could only rely on the haptic information about the object’s size and position provided by their left hand (Figure 2, fifth row). In the V_Full_ block, participants were provided with vision from the start and for the whole duration of each trial, whereas in the HV_Full_ block both visual and haptic information of the target object were simultaneously available for the whole duration of each trial (Figure 2, second last and last rows). The time interval from when the left hand was placed on the object to when the start tone was delivered was approximately 1 second long in the H_Full_, HV_Full_ and HV_t_ blocks.

The two sessions were performed in sequence, with the HV_t_ and the V_t_ blocks always preceding the other blocks, whereas the blocks were randomized within each session. The sequence containing the HV_t_ and the V_t_ blocks was always run first to prevent any learning effects due to full vision of the hand trajectory in the HV_Full_ and V_Full_ blocks. The two object positions and the visual exposure durations in the HV_t_ and V_t_ blocks were randomized within each block. Each of the eleven conditions (HV_50_, HV_100_, HV_200_, HV_400_, V_50_, V_100_, V_200_, V_400_, HV_Full_, V_Full_ and H_Full_) included 20 trials for the 300 mm position and 10 trials for the 150 mm position for a total of 330 trials per participant. To become accustomed to the task and the conditions, participants completed a training session in which ten trials were run with the object positioned at 300 mm before each block.

### 2.4 Data analysis

Kinematic data were analyzed in R (R Core Team, 2020). The raw data were smoothed and differentiated with a third-order Savitzky-Golay filter with a window size of 21 points. These filtered data were then used to compute velocities and accelerations of the index finger in three-dimensional space. Movement onset was defined as the moment of the lowest, non-repeating index finger acceleration value prior to the continuously increasing index acceleration values (Volcic and Domini, 2016), while the end of the movement was defined by applying the same algorithm, but starting from the end of the recorded values. From the 2860 trials directed toward the 300 m position we discarded from further analysis the trials in which the end of the movement was not captured correctly or in which the missing marker samples could not be reconstructed using interpolation. The exclusion of these trials (153 trials, 5.3% of all trials) left us with 2708 trials for the final analysis. For each trial we calculated the endpoint error along the azimuth and depth directions defined as the signed distance from the center of the object. In this way, positive values in depth and azimuth corresponded to an overshoot and a rightward displacement with respect to the center of the object, respectively.

To define at which stage of the movement vision was withheld in the HV_t_ and V_t_ blocks, we have calculated the index finger position (and its standard deviation) when the goggles turned opaque. We found that in all the 50 ms and 100 ms conditions the hand did not start moving yet as it still was at the home position (0 ± 0.1 mm of displacement), while in the HV_200_ and V_200_ conditions the index finger was 2 ± 5 mm and 0.3 ± 1 mm from the home position, respectively. In the HV_400_ condition the index finger was 135 ± 66 mm from the home position, whereas in the V_400_ condition it was 93 ± 66 mm from the home position. Therefore, in the 50 ms, 100 ms and 200 ms conditions vision was delivered only during the movement planning phase and very early movement execution, whereas in the 400 ms conditions participants were able to see the hand for the first ∼30% of the movement trajectory before vision was withheld.

### 2.5 Statistical analysis

Our main variables of interest were the within-participant endpoint variabilities in the azimuth and depth directions. We estimated these variabilities with a multivariate Bayesian linear mixed-effects model using the brms package (Bürkner et al, 2017), which implements Bayesian multilevel models in R using the probabilistic programming language Stan (Carpenter et al, 2017). The multivariate model used to fit the endpoint errors in the azimuth and depth directions included as the fixed-effect (predictor) the categorical variable Condition (HV_50_, HV_100_, HV_200_, HV_400_, V_50_, V_100_, V_200_, V_400_, HV_Full_, V_Full_ and H_Full_). The estimates of the Condition parameters (β_*Condition*_) thus correspond to the average endpoint error along the azimuth and depth direction of each Condition, whereas the σ_*Condition*_ parameters correspond to the within-participant endpoint variability in azimuth and depth directions.

The model included the independent random (group-level) effects for subjects. The model was fitted considering weakly informative prior distributions for each parameter to provide information about their plausible scale. We used Gaussian priors for the Condition fixed-effect predictor (β_*Condition*_: mean = 0 and sd = 20, σ_*Condition*_: mean = 0 and sd = 15), whereas for the group-level standard deviation parameters and sigmas we used zero-centered Student *t* -distribution priors (*df* = 3, scale = 10). Finally, we set a prior over the correlation and residual correlation matrix that assumes that smaller correlations are slightly more likely than larger ones (LKJ prior set to 2).

We ran four Markov chains simultaneously, each for 4,000 iterations (1,000 warm-up samples to tune the MCMC sampler) with the delta parameter set to 0.9 for a total of 12,000 post-warm-up samples. Chain convergence was assessed using the 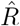 statistic (all values equal to 1) and visual inspection of the chain traces. Additionally, predictive precision of the fitted models was estimated with leave-one-out cross-validation by using the Pareto Smoothed importance Sampling (PSIS). All Pareto k values were below 0.5.

The posterior distributions we have obtained represent the probabilities of the parameters conditional on the priors, model and data, and, they represent our belief that the “true” parameter lies within some interval with a given probability. We summarized the posterior distributions related to the estimated σ_*Condition*_ (i.e., endpoint variability along the azimuth and depth directions) by computing the mean and the 95% credible intervals. The 95% credible interval specifies the interval that includes with a 95% probability the true value of a specific parameter. To evaluate the differences between parameters of a pair of conditions, we have subtracted the posterior distributions of *σ*_*Condition*_ weights between the conditions of interest. The resulting distributions are denoted as the credible difference distributions and are again summarized by computing the mean and the 95% credible intervals. For statistical inferences about the σ_*Condition*_ parameters we assessed the overlap of the 95% credible intervals with zero. A 95% credible interval that does not span zero is taken as evidence that the model parameters in the two conditions differ from each other.

## 3 Results

The conditions in which vision was constantly available (V_Full_ and HV_Full_) showed the lowest endpoint variability (Figure 3). Interestingly, the endpoint variabilities in the HV_t_ and V_t_ blocks are clustered in two different regions of the plane defined by the variabilities in the azimuth and depth directions. The endpoint variabilities of the V_t_ block are positioned in the top-right region of the plane, whereas the endpoint variabilities of the HV_t_ block lay in between the HV_Full_ and H_Full_ conditions, spreading along the diagonal line that indicates equal variability in azimuth and depth. Notably, the endpoint variabilities in the HV_t_ block are lower than the endpoint variabilities in the H_Full_ condition in both azimuth and depth directions suggesting that a brief glimpse at the haptic target is sufficient to trigger multisensory integration that influences movement execution.

**Figure 3:**
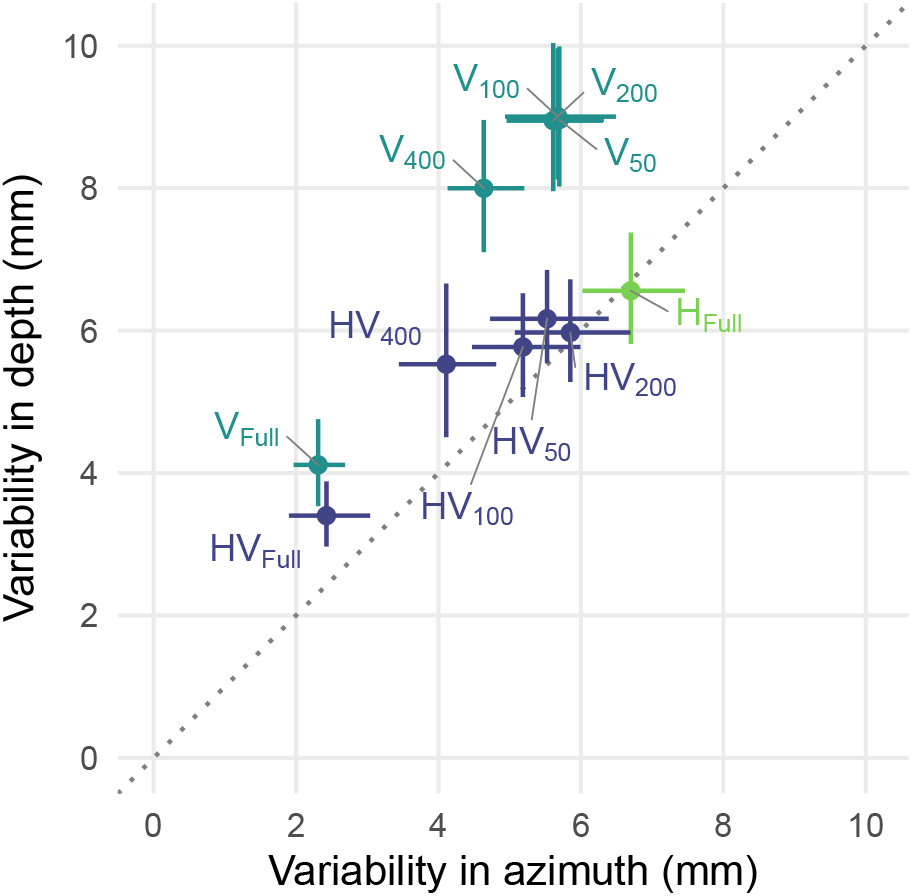
Estimates of the endpoint variability in the azimuth and depth directions of all the conditions. The dots represent the mean and the error bars denote the 95% credible intervals of the estimates. The grey dashed line represents the points of equal variability in the depth and azimuth directions

To thoroughly explore how vision and haptics are combined during movements guided by both modalities, pairwise comparisons between conditions were performed. Specifically, we compared: 1) the unisensory (H_Full_ and V_Full_) and multisensory (HV_Full_) conditions to define how the endpoint variability is modulated according to the available sensory information; 2) the conditions within each V_t_ and HV_t_ blocks to determine how endpoint variability is affected by changes in visual exposure; 3) the conditions of the HV_t_ block with the conditions of the V_t_ and the H_Full_ blocks to establish how effective and rapid visuo-haptic integration is; 4) the observed and the predicted endpoint variabilities to verify if the changes in precision in the multisensory conditions adhere to the maximum-likelihood model of sensory integration that combines the unisensory conditions.

### 3.1 Endpoint variability decreases in multisensory reaching

To determine how action performance is modulated according to the available sensory information, we compared the endpoint variability between the H_Full_, V_Full_, and HV_Full_ conditions. The comparison between V_Full_ and H_Full_ revealed an advantage of vision over haptics in both the depth and azimuth directions (Figure 4, left panel). The comparison between HV_Full_-H_Full_ and HV_Full_-V_Full_ conditions revealed an overall lower endpoint variability in the multisensory condition compared to each unisensory condition (Figure 4, middle and right panels). Interestingly, this advantage was direction-specific, evidencing a different contribution of haptics and vision within the multisensory integration process. On one hand, the comparison between the HV_Full_ and H_Full_ conditions showed that adding visual information decreased endpoint variability in both the depth and azimuth directions (Figure 4, central panel). On the other hand, the comparison between the HV_Full_ and the V_Full_ conditions showed that adding haptic information decreased the endpoint variability only in the depth direction (Figure 4, right panel). These results corroborate the finding that, in multisensory conditions, haptics mainly improves action precision along the depth dimension, whereas vision improves it along the azimuth direction (van Beers et al, 2002; Monaco et al, 2010).

**Figure 4:**
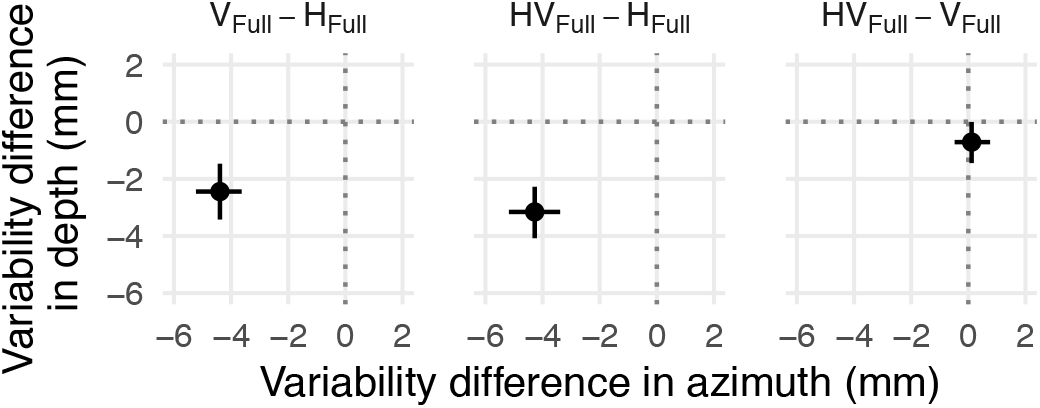
Credible difference distributions of the comparisons between the H_Full_, V_Full_ and HV_Full_ conditions for the endpoint variability in the azimuth and depth direction. Dots represent the mean and the error bars represent the 95% credible intervals of the credible difference distributions.

### 3.2 The effect of the visual exposure duration on endpoint variability

To determine how the changes in visual exposure duration affect endpoint variability we have sequentially compared the conditions within the V and HV blocks. Withholding vision during the initial stages of the movement (V_400_ and HV_400_) led to an increase in endpoint variability in both the azimuth and depth directions compared to when vision was available throughout the movement (V_Full_ and HV_Full_). Endpoint variability worsened whether haptics was available or not, but the increase was more pronounced when only vision was provided (Figure 5, first column, V_400_−V_Full_ and HV_400_−HV_Full_ comparisons). The further reduction in visual exposure duration (V_200_ and HV_200_), which limited visual availability to only the movement preparation phase, led to an additional increase in endpoint variability (Figure 5, second column, V_200_−V_400_ and HV_200_−HV_400_ comparisons). Whereas endpoint variability in V_200_ increased approximately equally in both the azimuth and depth directions, the increment in endpoint variability in HV_200_ was constrained to the azimuth direction. Interestingly, endpoint variability did not worsen when the visual exposure duration was further reduced from 200 ms to 100 ms (Figure 5, third column, V_100_−V_200_ and HV_100_−HV_200_ comparisons), and, from 100 ms to only 50 ms (Figure 5, fourth column, V_50_−V_100_ and HV_50_−HV_100_ comparisons).

**Figure 5:**
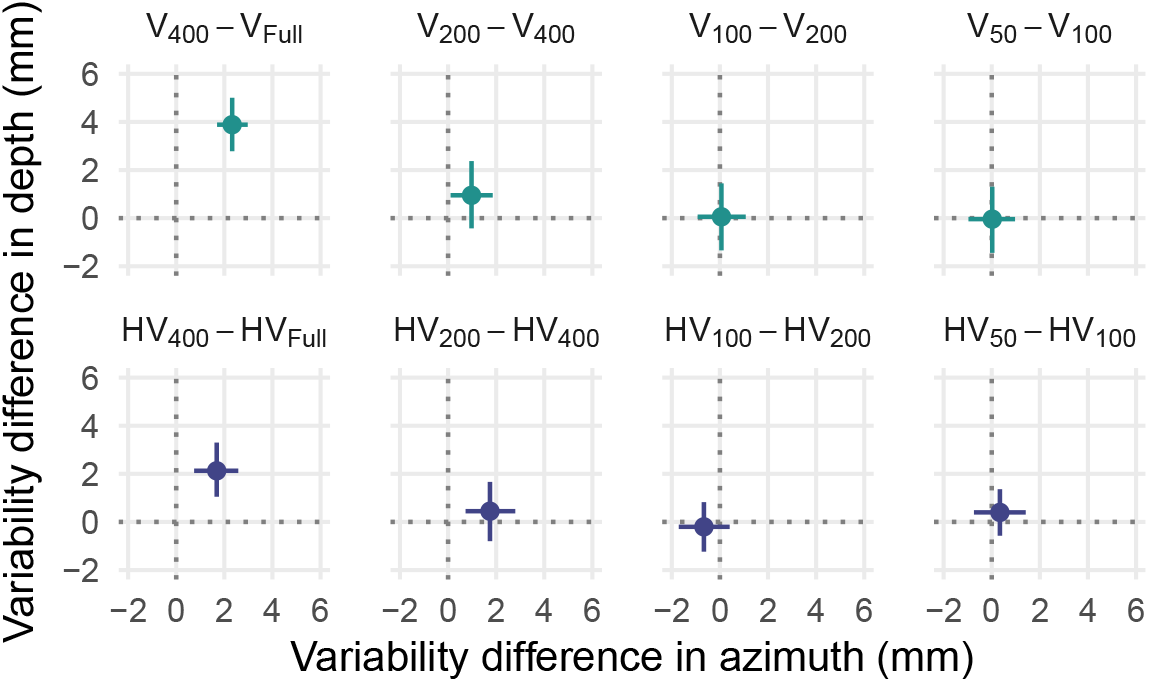
Top row: Credible difference distributions of the endpoint variability in the azimuth and depth direction for the comparisons between V_400_ and V_Full_ and between the conditions of the V_t_ block. Bottom row: Credible difference distributions of the endpoint variability in the azimuth and depth direction for the comparisons between HV_400_ and HV_Full_ and between the conditions of the HV_t_ block. Dots represent the mean and the error bars represent the 95% credible intervals of the credible difference distributions.

These findings clearly show that action precision depends on the availability of visual information during movement execution. However, the more moderate decrease in precision in the HV_t_ conditions compared to the V_t_ conditions also shows that haptic information can partially compensate for the lack of vision.

### 3.3 A brief visual exposure is sufficient for improved endpoint variability

To determine whether action performance in HV_t_ is a product of multisensory integration and it is thus better than the performance when only the single senses are available, we further contrasted the conditions of the HV_t_ block with the H_Full_ block and with the conditions of the V_t_ block. Whereas the conditions of the HV_t_ and the H_Full_ blocks share the same haptic information and they only differ in terms of visual information, the conditions of the HV_t_ and the V_t_ blocks provide the same visual information, but they differ with regard to the presence/absence of haptic information. Thus, the HV−H and the HV−V contrasts can provide an answer about how does action performance improve when both vision and haptics are available.

We found that the endpoint variability in all the conditions of the HV_t_ block was reduced compared to the H_Full_ condition (with the exception of the HV_200_−H_Full_ comparison) and of the V_t_ block (Figure 6), providing further evidence of multisensory integration. The multisensory advantage was manifest at all visual exposure durations, even when vision was provided for only 50 ms before movement onset (Figure 6, rightmost column), which is remarkable considering that 50 ms are less than the time of an eye blink (Garten, 1898; Weiss, 1911).

**Figure 6:**
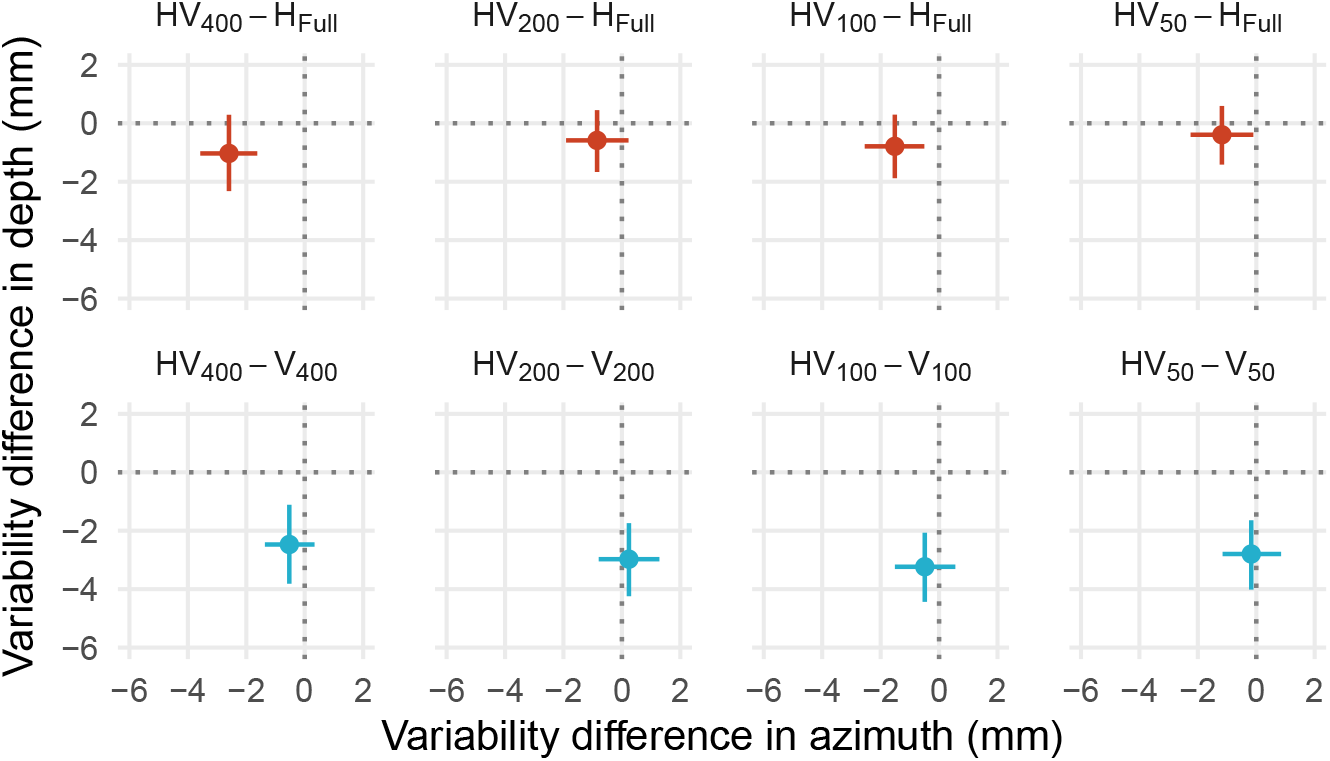
Top row: Credible difference distributions of the endpoint variability in the azimuth and depth direction for the comparisons between the H_Full_ condition and the conditions of the HV_t_ block. Bottom row: Credible difference distributions of the endpoint variability in the azimuth and depth direction for the comparisons between the conditions of the V_t_ and H_Vt_ blocks. Dots represent the mean and the error bars represent the 95% credible intervals of the credible difference distributions.

Interestingly, these multisensory advantages were, again, direction-specific. The comparisons between the H_Full_ block and the conditions of the HV_t_ block revealed that, when visual on-line control was not available, a brief visual exposure decreased endpoint variability mainly along the azimuth direction but not along the depth direction (Figure 6, top row). On the contrary, the comparisons between the conditions of the HV_t_ and the V_t_ blocks showed that the availability of haptic information reduced endpoint variability along the depth direction but not along the azimuth direction (Figure 6, bottom row).

It is possible, however, that the differences in precision among all the conditions that rely on haptic information were not due to the visual exposure durations, but rather to the decay of proprioceptive information (proprioceptive drift). According to this alternative interpretation, precision should degrade more the more time has passed since the left hand has been placed around the target object, because the proprioceptive information about the target position decays over time (Cameron et al, 2015; Goettker et al, 2020). This decay could be limited to the planning phase or it could extend over both the planning and execution phases. Thus, if a link exists between proprioceptive decay and precision, we should find that the less precise conditions have either longer reaction times (time from the start sound to the movement onset spanning the whole planning phase) or longer trial times (time from the start sound to the movement end spanning both planning and execution phases). To compare the reaction time and the trial time durations among conditions we ran two additional Bayesian linear mixed-effects models by considering the Condition as the fixed-effect predictor (for details, see Supplementary Material). We found no differences in both reaction time and trial time among all the conditions that relied on haptic information, except for the reaction time in the HV_Full_ condition which was slightly shorter than in the HV_400_ condition (Figure 7). These results rule out the alternative explanation that the differences in precision were due to proprioceptive decay, which presumably occurs over longer time intervals spanning multiple trials (Wann and Ibrahim, 1992; Desmurget et al, 2000; Smeets et al, 2006).

**Figure 7:**
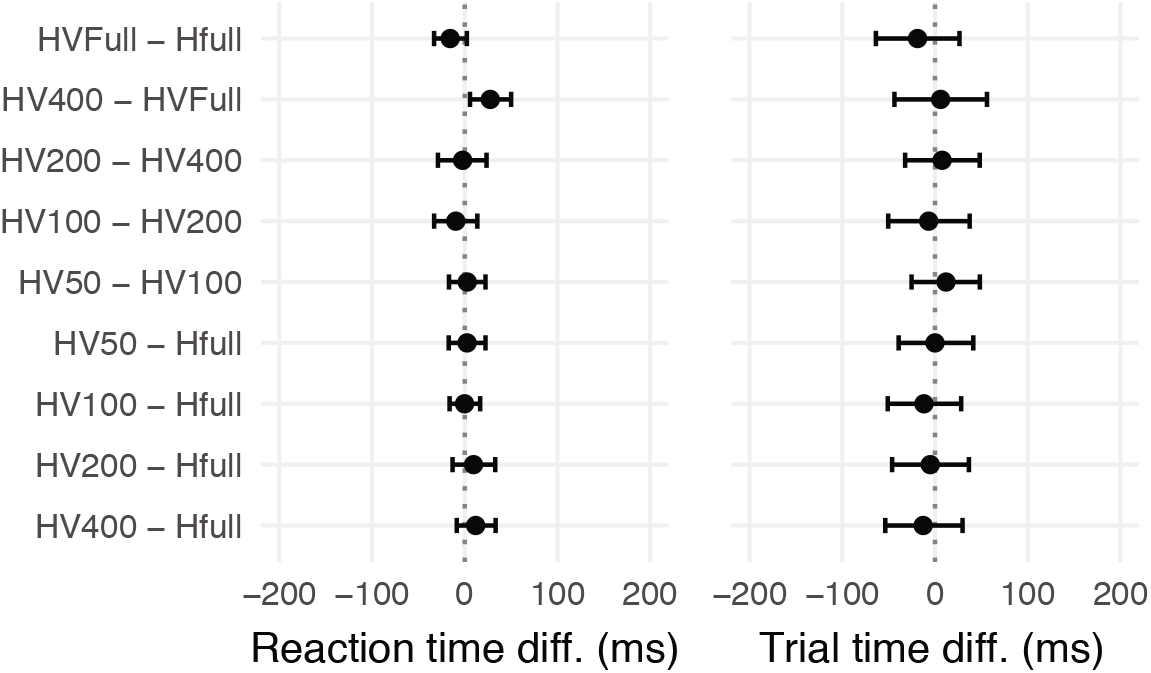
Credible difference distributions of the reaction time (left column) and the trial time durations (right column). Dots represent the mean and the error bars represent the 95% credible intervals of the credible difference distributions.

### 3.4 Optimal and near-optimal integration of haptic and visual inputs

If the independent haptic and visual inputs are optimally combined, we should have observed a reduction of endpoint variability consistent with the predictions made by a maximum-likelihood model of sensory integration of these unimodal inputs. The combined estimates should have maximal precision (i.e., minimum variance) and they should be more precise than either the haptic or visual estimates alone (Cochran, 1937; Ghahramani et al, 1997; Ernst and Banks, 2002). Specifically, the predicted endpoint variability in each HV condition (HV_Full_, HV_400_, HV_200_, HV_100_, HV_50_) should be equal to:

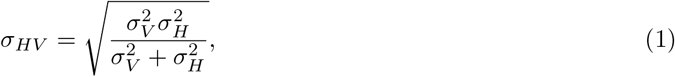

where *σ*_*H*_ is the endpoint variability in the H_Full_ condition and *σ*_*V*_ is the endpoint variability in the respective V condition (V_Full_, V_400_, V_200_, V_100_, V_50_).

In the previous sections we have shown that the precision in all multisensory conditions was highest than the highest precision of the unisensory conditions, either in the azimuth or in the depth directions, which is an important indication of multisensory integration. However, optimal integration was not achieved in all conditions. The predicted endpoint variability was indistinguishable from the observed endpoint variability only in the HV_Full_ and HV_400_ conditions (Figure 8, first and second columns). Instead, the predicted endpoint variability was smaller than the observed endpoint variability in the azimuth direction for the HV_200_ and HV_100_ conditions, and, in both the azimuth and the depth directions for the HV_50_ condition (Figure 8, third, fourth and fifth columns), suggesting near-optimal integration (Rohde et al, 2016).

**Figure 8:**
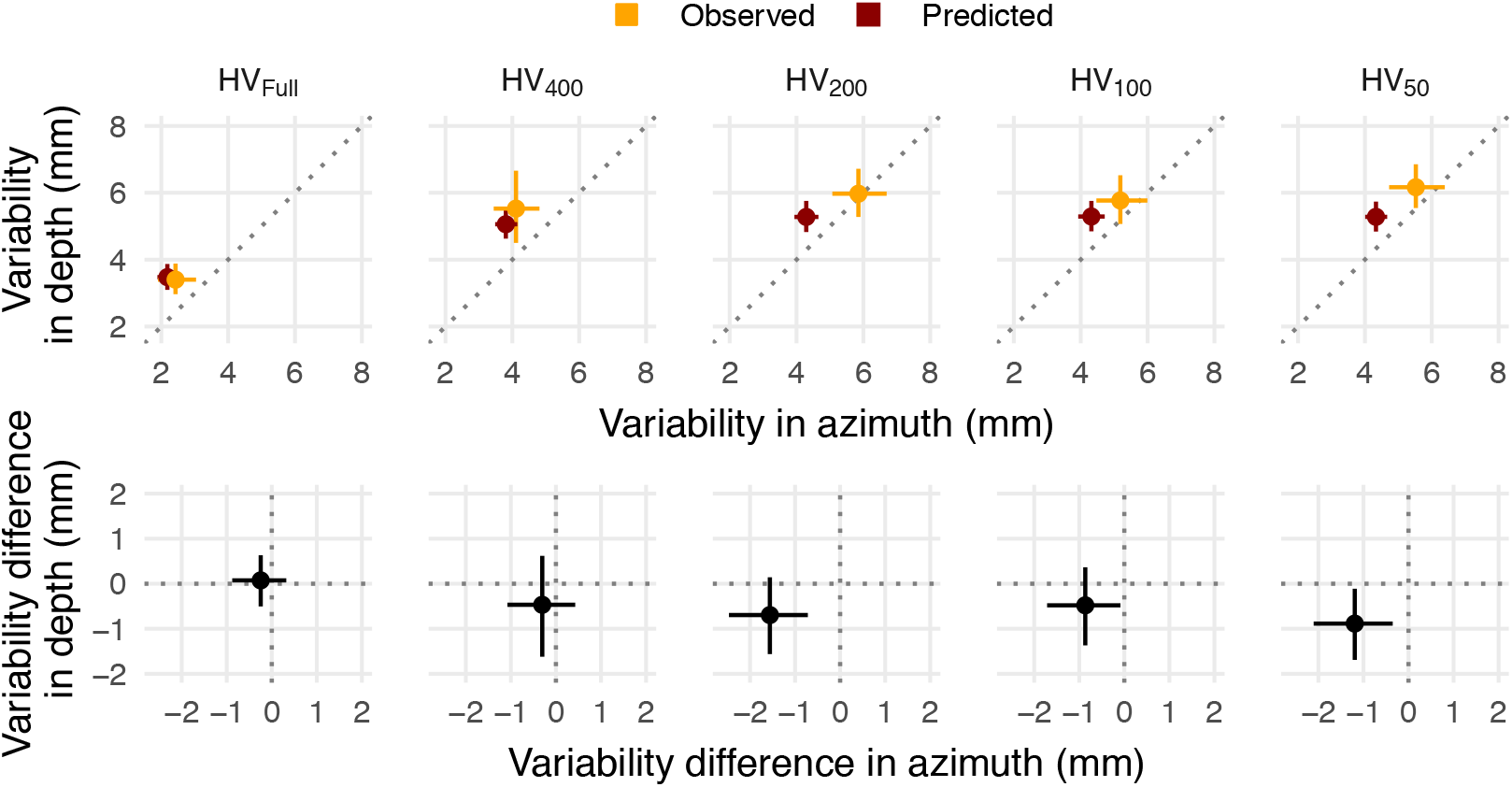
Top row: Predicted and observed endpoint variabilities in the azimuth and depth directions. Bottom row: Differences between the predicted and the observed endpoint variabilities in the azimuth and depth directions. Dots represent the mean and the error bars represent the 95% credible intervals of the credible difference distributions.

## 4 Discussion

This study shows that reaching movements towards visuo-haptic targets are guided by a multisensory integration process even when the visual exposure to the haptic target during the planning phase is as short as 50 ms. Supplementing haptic information with very brief vision is thus sufficient to trigger a multisensory advantage that persists until the end of the action, even though vision provided only during the planning phase does not lead to the same level of precision as when it is provided during the whole movement (Keele and Posner, 1968; Elliott and Madalena, 1987; Blouin et al, 1993; Rossetti et al, 1994; Khan et al, 2002, 2006; Kennedy et al, 2015; Tremblay et al, 2017; Desmurget et al, 1997; Monaco et al, 2010; Cameron and López-Moliner, 2015) In these conditions, a superior action performance is observed compared to when only haptic information is accessible throughout the whole movement (H_Full_) or only vision is provided during the planning phase (V_t_). We therefore conclude that, when visual availability is minimal (HV_t_), the precision of reaching movements is mostly determined by the older, but most reliable, visuo-haptic information rather than by the more recent, but the less reliable sense.

Even though all visual exposure durations improved precision in multisensory conditions, the longer visual exposure duration (400 ms) led to higher precision than the shorter durations, which were instead all comparably precise. This difference could be attributed to the fact that participants simply had more time to plan the movement, or, to the fact that the longest visual exposure duration included both the planning phase and a short initial part of the movement, whereas the shorter durations were limited only to the planning phase. The latter interpretation seems more likely, since seeing the reaching hand at the initial stage of the action has been, indeed, shown to be essential to specify the initial movement direction and thus affect the final endpoint precision (Sainburg et al, 2003; Sarlegna and Sainburg, 2007; Bagesteiro et al, 2006).

Most interestingly, our study provides clear evidence that it takes no more than 50 ms of vision to successfully start the visuo-haptic integration process that increases the endpoint precision. We can exclude that this improvements were due to a learning effect, since the visual exposure durations were randomized within the HV_t_ block, and, this block was performed before any block that provided vision during the movement. And, we can exclude that the differences in precision were caused by the decay of proprioceptive information, since the reaction time and trial time durations were very similar among the conditions that relied on haptic information. Thus, a sufficiently rich representation of the scene and the to-be-reached object can be extracted in less than the duration of an eye blink (Garten, 1898; Weiss, 1911). This ability could be very advantageous especially when moving our hands in cluttered environments in which a brief look toward the target object would be sufficient to increase the precision of a reaching movement, without limiting vision to promptly move to the next object of interest (Johansson et al, 2001).

The current findings also show several similarities with previous studies on multisensory reaching. First, we confirmed that actions guided by simultaneous visual and haptic inputs available throughout the whole movement are more precise than actions guided by visual only inputs, which were, in turn, more precise than actions guided by haptic only inputs (van Beers et al, 1996). Second, we confirmed the direction-specific contribution of vision and haptics in multisensory reaching. Whereas vision improved precision along the azimuth direction, haptics played a role in increasing precision in the depth direction (van Beers et al, 1999b,a). Interestingly, this direction-specificity persisted also in the HV_t_ conditions in which multisensory information was available only during the planning phase (Monaco et al, 2010; Cameron and López-Moliner, 2015).

Lastly, we showed that haptics and vision were optimally integrated, in accordance with a maximum-likelihood model of sensory integration, only when vision was provided during the whole movement execution or at its early stage (i.e., HV_Full_ and HV_400_ conditions, respectively). A near-optimal integration was, instead, observed when vision was only briefly provided during the planning phase. Failing to achieve optimal integration also in these conditions might have been due to several reasons. One explanation could be that reaching movements were guided only by haptic information in a small proportion of trials, because the visual input was missed in those occasions (e.g., if an eye blink occurred exactly in the moment vision was provided). An alternative but equally plausible explanation could be that the output of the multisensory integration process might have slowly decayed over time (Westwood et al, 2003).

Taken together, our results show that 50 ms of vision during the action planning phase are sufficient to trigger a direction-specific multisensory integration process that increases the precision of reaching actions, in line with optimal integration models. Our study suggests that the oldest, but most reliable, multisensory information determines action precision, even when a less reliable unisensory information is available until the end of the movement.

## CRediT author statement

**Ivan Camponogara:** Conceptualization, Methodology, Software, Validation, Formal analysis, Investigation, Data Curation, Writing - Original Draft, Writing - Review & Editing, Visualization. **Robert Volcic:** Conceptualization, Methodology, Software, Validation, Formal analysis, Data Curation, Resources, Writing - Review & Editing, Visualization, Supervision.

## Additional information

### Competing interests

The authors declare no competing interests.

Correspondence and requests for materials should be addressed to I.C.

## Notes

### Competing Interest Statement

The authors have declared no competing interest.

